# Autophagy regulates the localization and degradation of p16^INK4a^

**DOI:** 10.1101/521682

**Authors:** Philip R. Coryell, Supreet K. Goraya, Katherine A. Griffin, Margaret A. Redick, Samuel R. Sisk, Jeremy E. Purvis

**Affiliations:** Department of Genetics, University of North Carolina at Chapel Hill, Chapel Hill, NC 27599-7264; Curriculum for Bioinformatics and Computational Biology, University of North Carolina at Chapel Hill, Chapel Hill, NC 27599-7264; Lineberger Comprehensive Cancer Center, University of North Carolina at Chapel Hill, Chapel Hill, NC 27599-7264; Computational Medicine Program, University of North Carolina at Chapel Hill, Chapel Hill, NC 27599-7264

## Abstract

The tumor suppressor protein p16^INK4a^ (p16) is a well-established hallmark of aging that induces cellular senescence in response to stress. Previous studies have focused primarily on p16 regulation at the transcriptional level; comparatively little is known about the protein’s intracellular localization and degradation. The autophagy-lysosomal pathway has been implicated in the subcellular trafficking and turnover of various stress-response proteins, but it is unclear whether p16 is involved in these pathways. Here, we investigate the role of autophagy, vesicular trafficking, and lysosomal degradation on p16 expression and localization in human epithelial cells. Time-lapse fluorescence microscopy using an endogenous p16-mCherry reporter revealed that autophagy induced by genotoxic stress stimulates rapid p16 recruitment to acidic cytoplasmic vesicles. When vesicular acidification was inhibited by NH_4_Cl, nuclear p16 levels increased. Single-cell imaging revealed that p16 localizes to lysosomes upon stress, implicating the autophagy pathway as a regulator of p16 localization. Blocking autophagy with bafilomycin, chloroquine, or NH_4_Cl resulted in elevated p16 protein levels without increased transcription. Increased p16 coincided with accumulation of autophagosome chaperone p62/SQSTM1 (p62) and decreased levels of phosphorylated-Rb. Furthermore, chloroquine caused p16 aggregation within stalled vesicles containing autophagosome marker LC3, demonstrating that p16 is transported and degraded by the autophagy-lysosomal pathway. Knockdown of p62 resulted in delocalization of p16 aggregates to autophagosomes, suggesting that p16 is targeted to these vesicles by p62. Taken together, these results implicate the autophagy pathway as a novel regulator of p16 degradation and localization, which could play a role in the etiology of cancer and age-related diseases.

## Introduction

The tumor suppressor protein p16^INK4a^ (CDKN2A, p16) is a member of the INK4 family of cyclin-dependent kinase inhibitors, which play a critical role in cell cycle regulation. Expression of p16 prevents cellular proliferation by binding and inhibiting cyclin-dependent kinases 4 and 6 (CDK4/6). In response to oncogene expression and prolonged DNA damage, p16 induces cellular senescence (permanent cell cycle arrest) [1]. As an organism ages, p16 accumulates in tissues, which triggers cellular senescence. Clearance of p16 expressing senescent cells has been linked to an increase in lifespan and a decrease in tumorigenesis [2]. The correlation between p16 expression and aging is so strong that p16 is commonly used as a biomarker for aging [3]. While the mechanisms regulating transcription of p16 have been well described, studies about the localization and degradation of the p16 protein are lacking.

p16 is expressed in both the nucleus and the cytoplasm [4];[5]. Whereas the role of p16 in the nucleus as an inhibitor of CDK4/6 is well understood, its subcellular localization and function in the cytoplasm remains mysterious. Immunohistological studies of patient tumors have suggested p16 localization as a possible indicator of clinical prognosis. However, many of these studies present contradictory claims that indicate a complex role for p16 localization in tumor progression. For example, cytoplasmic p16 has been reported to be a predictor of poor prognosis in patients with astrocytic brain tumors [6]. However, cytoplasmic p16 has also been reported as correlating with the absence of metastasis in other cancer types, such as melanoma [7]. Commonly used chemotherapeutic drugs such as etoposide and doxorubicin can increase p16 protein levels and induce senescence [8], but whether and to what extent these agents affect p16 localization has not been fully explored. Interestingly, p16 does not have a known nuclear localization signal (NLS) or a nuclear export signal (NES) [9], suggesting that an indirect mechanism of intracellular transport is responsible for shuttling p16 between different cellular compartments [10].

One potential mechanism for regulation of p16 localization is vesicular trafficking via the lysosomal endomembrane system. Lysosomes are cytoplasmic organelles involved in autophagy-mediated protein degradation. Like p16, lysosomes are involved in senescence-associated signaling pathways, and lysosome dysfunction has been linked to a myriad of age-related pathologies and a decrease in lifespan [11];[12];[13]. Similarly, lysosomes have also been targeted for lifespan extension therapies, such as intervention with rapamycin and spermidine [11]. Recent studies have expanded beyond protein degradation and explored the role of lysosomes in subcellular localization of stress-response proteins and the regulation of cell fate. For example, the mechanistic target of rapamycin (mTOR) was found to not only be recruited and degraded by lysosomes, but also play an important role in lysosome formation and regulation of the entire autophagy pathway [14]. Given the correlation of both autophagy and p16 expression with cellular aging and senescence, an intriguing hypothesis is that p16 localization, degradation, and regulation may be mediated by lysosomes and other members of this pathway. Previous experiments have shown that p16 can be degraded by the proteasome [15]; however, no literature exists to support whether regulation can also occur through other known degradation mechanisms such as the autophagy/lysosomal pathway.

As shown in **Figure 1**, the autophagy pathway consists of several sequential steps, beginning with stimulation by nutrient starvation or stress, followed by interaction between several endomembrane vesicles, and ending with the lysosomal degradation of proteins. Autophagy can be both selective and non-selective. For selective autophagy, ubiquitinated proteins or protein aggregates can be targeted for lysosomal degradation by ubiquitin-binding protein p62 (also known as sequestosome 1; SQSTM1). Although p16 does not contain a lysine residue, N-terminal ubiquitination of p16 has been reported [15]. p62-bound proteins are enveloped by autophagosomes, which are identifiable by autophagosome marker Membrane-bound microtubule-associated protein 1A/1B-light chain 3 (LC3). Autophagosomes then fuse to lysosomes (identifiable by Lysosomal-associated membrane protein 1 LAMP-1) containing low-pH dependent hydrolases, forming autolysosomes. The pH gradient of these vesicles lowers throughout this process, provoking the degradation of proteins within the autolysosome, including p62 and LC3. Autophagic flux, or the rate at which proteins are degraded by this pathway, can change in response to cellular stress and nutrient availability. Changes in autophagic flux can rapidly affect the localization and expression dynamics of proteins involved in this pathway [16]. Accordingly, researchers studying autophagy often employ inhibitors to capture proteins in transit within this pathway. Examples of well-characterized autophagy inhibitors include bafilomycin A1, chloroquine, and ammonium chloride, which act by preventing the fusion of autophagosomes and lysosomes, and by raising vesicular pH, which prevents hydrolase activity [17].

**Figure 1.**
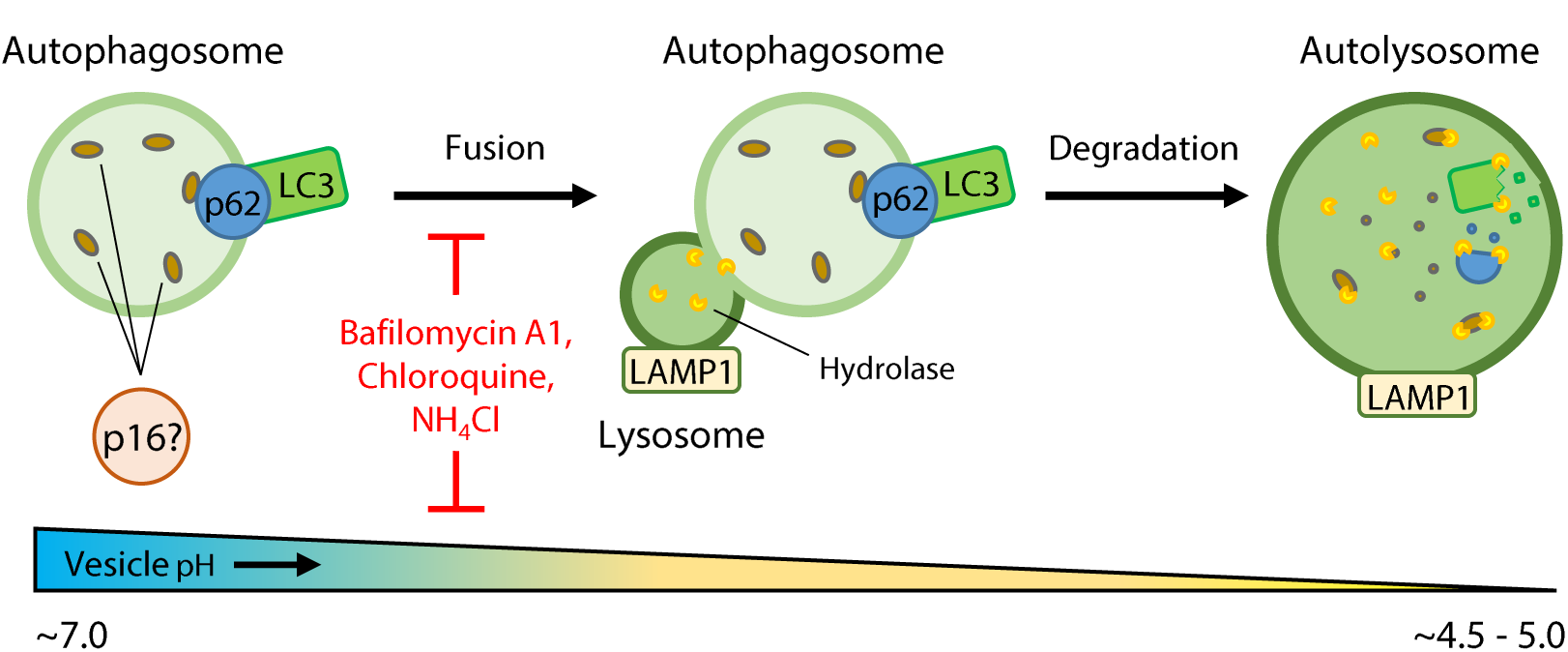
Autophagy pathway model depicting molecular markers and inhibitors. Autophagy chaperone protein p62 targets proteins destined for lysosome-mediated degradation to autophagosomes, which are identifiable by autophagasome membrane marker LC3. Autophagosomes fuse to lysosomes (identifiable by lysosome membrane marker LAMP-1) containing low-pH dependent hydrolases, forming autolysosomes. The pH gradient of these vesicles lowers throughout this process, provoking the degradation of proteins within the autolysosome, including p62 and LC3. Autophagy inhibitors bafilomycin A1, chloroquine, and ammonium chloride act by preventing the fusion of autophagosomes and lysosomes, and by raising vesicular pH, which prevents hydrolase activity.

In this study, we investigated the relationship between p16 and the autophagy/lysosomal pathway in human cells. By engineering a live-cell reporter for p16, we found that activation of autophagy caused p16 to accumulate in acidic cytoplasmic vesicles that stained positive for the lysosome marker LAMP-1. Neutralizing lysosomes led to increased p16 levels and preferential accumulation of nuclear p16. Chemical inhibition of autophagy led to increases in p16 protein as well as the autophagosome chaperone p62. Depletion of p62 abolished the ability of p16 to colocalize with autophagosomes. Taken together, these results show that p16 is localized and degraded through the autophagy/lysosomal pathway, implicating the autophagy pathway as a potential regulator of p16-induced senescence.

## Results

### Localization dynamics of p16 in response to autophagy and genotoxic stress

Autophagy is a highly dynamic process involving rapid protein transport and turnover known as autophagic flux. As a consequence, many autophagy markers and proteins targeted for degradation are difficult to measure [16, 18]. Immunostaining of fixed cells can capture protein localization only at a single point in time. Furthermore, permeabilization using common detergents, such as Triton X-100, can destroy membrane-bound organelles such as endosomes, lysosomes, and autophagosomes [19]. As an alternative approach, fluorescently tagged protein reporters have been employed to accurately visualize and track temporal changes of members of the autophagy pathway and proteins destined to this pathway for degradation [16, 18].

Taking this approach, we developed a live-cell reporter to monitor p16 protein expression in real time. A fluorescent p16-mCherry fusion protein was introduced at the endogenous p16 locus in human retinal pigment epithelial cells (RPE-1) cells using CRISPR-mediated homologous recombination. A p16-mCherry donor cassette for targeted homologous recombination was transfected into RPE-1 cells along with a Cas9 nuclease guided to an untranslated region near the p16 stop codon (**Supplemental figure 1A**). mCherry was selected because of its pH stability and ability to maintain fluorescence under acidic conditions, including within the lysosomal lumen [20, 21]. In order to differentiate nuclear from cytoplasmic p16-mCherry and monitor genotoxic stress, we also incorporated a constitutively expressed 53BP1-mVenus reporter in these cells (**Supplemental figure 1B**). 53BP1 remains diffusely expressed throughout the nucleus and also localizes to sites of DNA double-stranded breaks [22]. This resulted in the creation of a combined p16-mCherry 53BP1-mVenus reporter cell line (henceforth referred to as RPE p16-mCherry) that could be utilized for subcellular segmentation during live-cell fluorescence experiments (**Supplemental figure 1C**).

We first asked how p16 expression and localization changes in response to autophagy and stress. To do this, we treated endogenously-tagged RPE p16-mCherry reporter cells with either DMSO or 20 µM etoposide. Etoposide is a type-II topoisomerase inhibitor that induces DNA double-stranded breaks and activates autophagy [23]. To monitor the activation of autophagy, cells were also treated with Lysotracker, a live-cell chemical stain for V-ATPase activity in acidified vesicles. DMSO-treated control cells exhibited sparse Lysotracker staining, as well as diffuse cytoplasmic and concentrated nuclear p16-mCherry (**Figure 2A**). In contrast, etoposide-treated cells displayed bright cytoplasmic puncta in response to Lysotracker, demonstrating that etoposide was sufficient to trigger autophagy. Moreover, these cells accumulated cytoplasmic p16-mCherry puncta that co-localized with Lysotracker, demonstrating that p16-mCherry localizes to acidic cytoplasmic compartments in response to etoposide. Time-lapse images revealed that p16-mCherry puncta began forming approximately 10-12 hours after etoposide treatment (**Figure 2B**).

**Figure 2.**
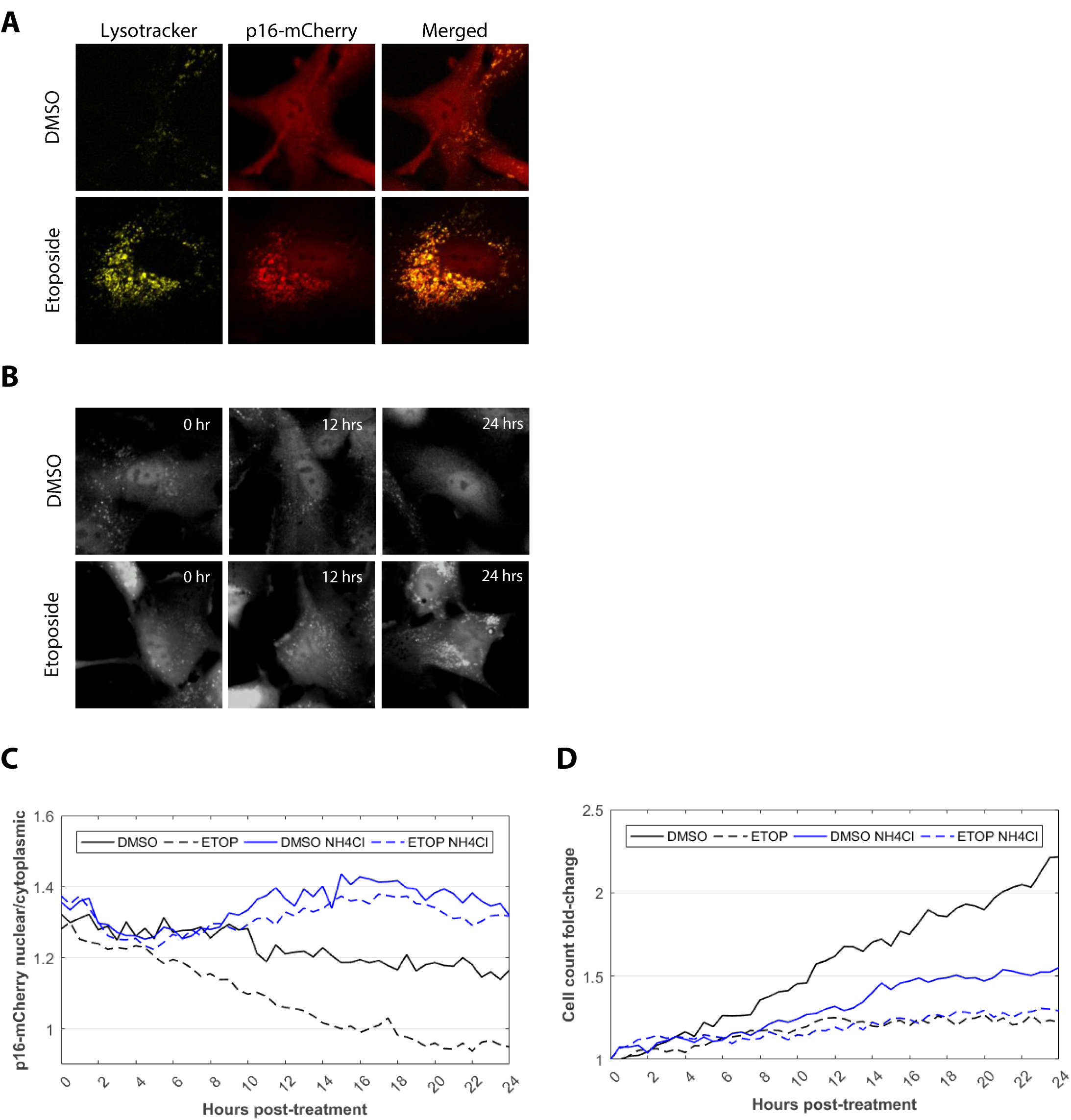
Dynamics of p16 localization in response to autophagy and stress. **A)** Human RPE-1 p16-mCherry cells 24 h after DMSO or etoposide treatment. p16-mCherry is shown in red; Lysotracker, which stains acidic organelles (pH ~ 4.5-5), is shown in yellow. **B)** Time-lapse images of p16-mCherry cells after DMSO or etoposide treatment. **C)** Ratio of nuclear to cytoplasmic p16-mCherry signal across cell population **D)** Cell counts for each treatment measured by the number of 53BP1-mVenus nuclei counted each frame. *n >* 100 cells per condition at each timepoint.

We next asked how disruption of autophagy affects the expression and subcellular localization dynamics of p16. Autophagy can be inhibited by exposing cells to ammonium chloride (NH_4_Cl), which raises the luminal pH of intracellular vesicles and prevents the activation of degradative enzymes inside lysosomes [24]. Treatment with 40 mM NH_4_Cl for 24 hours was sufficient to quench Lysotracker signal in etoposide-treated cells, confirming that vesicular acidification was inhibited by this chemical (data not shown). RPE p16-mCherry cells were treated with either DMSO or 20 µM etoposide alone, or in combination with 40 mM NH_4_Cl. We then performed time-lapse fluorescence microscopy for 24 h and compared the ratio of nuclear versus cytoplasmic p16-mCherry. In DMSO-only control cells, p16-mCherry remained nuclear concentrated for 24 h (**Figure 2C**). NH_4_Cl-inhibition of autophagy resulted in further nuclear p16-mCherry enrichment, diverging from the control population approximately seven hours after treatment. Etoposide shifted p16 enrichment to the cytoplasm, diverging from the control population at approximately 2.5 h after treatment. However, exposure to NH_4_Cl in etoposide-treated cells rescued this phenotype, enhancing nuclear enrichment of p16-mCherry similar to NH_4_Cl-treatment in control cells. To assess the effect of these treatments on the cell cycle, we also measured the change in cell population over the course of the experiment (**Figure 2D**). After 24 hours, the number of DMSO treated cells was doubled. DMSO with NH_4_Cl (which induced nuclear localization of p16-mCherry) perturbed the cell cycle, resulting in a 50% reduction in growth after 24 hours compared to DMSO alone. Etoposide (which caused cytoplasmic localization of p16-mCherry) induced cell cycle arrest, and resulted in only a 25% increase in cell population. Interestingly, etoposide with NH_4_Cl (which induced nuclear p16-mCherry) also resulted in a 25% increase, despite the divergent subcellular localization of p16-mCherry compared to etoposide-only. This suggests that the induction of cell-cycle arrest within 24 hours of etoposide treatment was not driven by changes in p16 localization.

Together, these data suggest that induction of autophagy sequesters p16 to acidic compartments in the cytoplasm and thereby attenuates its nuclear localization. Etoposide treatment further enhances cytoplasmic accumulation of p16 and induces cell cycle arrest. However, disruption of autophagy via NH_4_Cl reverses this trend by enriching p16 in the nucleus and preventing cytoplasmic build-up. These results implicate the autophagy pathway as a partial regulator of p16 protein localization.

### Lysosomes recruit p16 during autophagy and attenuate its nuclear localization during genotoxic stress

The autophagy-mediated protein degradation pathway involves the acidification of lysosomes in order to activate low-pH dependent hydrolases within the lysosomal lumen. Our live-cell experiments revealed co-localization between acidic organelles and p16-mCherry in response to stress. To confirm this finding in a non-reporter cell line, we developed an immunofluorescence protocol suitable for segmenting and quantifying p16 levels in lysosomes within single-cells (**Materials and Methods**). This protocol was specifically designed to avoid the destruction of membrane-bound organelles, like lysosomes, by permeabilizing fixed cells with digitonin, a selective detergent that punctures the plasma membrane while leaving endomembrane vesicles intact [25].

Using this modified immunofluorescence technique, we first tested for p16 and lysosome co-localization in unmodified RPE-1 cells undergoing autophagy and genotoxic stress. Control cells treated with DMSO displayed minor visual overlap between p16 and lysosomal-associated membrane protein 1 (LAMP-1) (**Figure 3A**). We then induced concurrent DNA damage and autophagy by treating cells with 20 µM etoposide for 72 h. Etoposide treatment resulted in easily visible p16 puncta that appeared interspersed among the cytoplasm. These puncta co-localized with LAMP-1, suggesting that a proportion of p16 was recruited to lysosomes upon activation of autophagy. Quantification of total p16 fluorescence intensity revealed that the amount of p16 within lysosomes was significantly increased by etoposide treatment (**Figure 3B**). After normalizing for changes in the size and number of lysosomes per cell, the mean p16 intensity per lysosome per cell was also increased after etoposide treatment (**Figure 3C**), suggesting active recruitment of p16 to lysosomes during autophagy.

**Figure 3.**
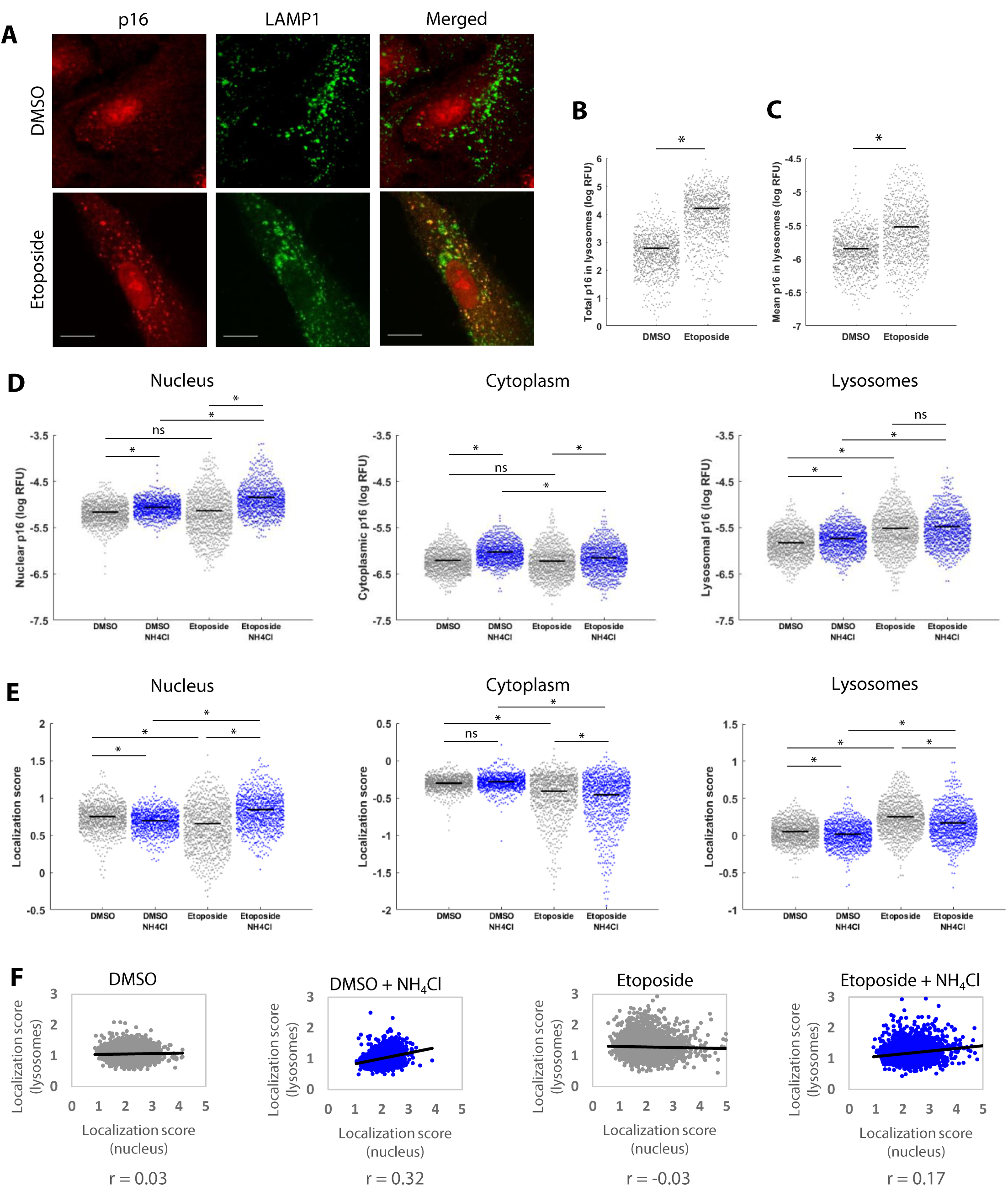
p16 localization in response to genotoxic stress and autophagy. RPE-1 cells were treated with either DMSO or 20 μΜ etoposide for 72 hours. For lysosome inhibition experiments, cells were also treated at 48 h with 40 mM NH_4_CI for an additional 24 h. Cells were then fixed for immunostaining and fluorescence quantification. For violin plots, each dot is one of 1000 cells randomly sampled from each treatment group. Blue dots indicate cells treated with NH_4_CI. RFU = relative fluorescence units. Black line = mean. p-values determined by ANOVA and multiple comparison analysis. * = p < 0.01. ns = p > 0.01. **A)** Immunofluorescence showing p16 (red) or Lysosomal-associated membrane protein 1 (LAMPI, green). Scale bars = 25 μm. **B)** Comparison of summed p16 pixel intensities overlapping LAMP1 in each cell. **C)** Sum of all p16 pixel intensities overlapping LAMP1 normalized to the number and size of lysosomes in each cell. **D)** Mean p16 within each subcellular compartment per cell. **E)** p16 localization score for each subcellular compartment per cell. Localization score = mean p16 of compartment divided by mean whole cell p16. **F)** Regression plots comparing p16 lysosomal localization score vs. p16 nuclear localization score. r = Pearson's correlation coefficient.

Previous studies have demonstrated considerable cross-talk between subcellular compartments, including between lysosomes and the nucleus [26, 27]. Therefore, we quantified the effects of genotoxic stress and lysosome dysfunction on p16 localization to three subcellular compartments within individual cells: cytoplasm, nucleus, and lysosomes. To do this, we first segmented the nuclei and lysosomes of all cells using chemical and antibody staining of the nucleus (DAPI) and lysosomes (LAMP-1), respectively. Next, using a highly amplified image of p16 antibody staining, we located the exterior borders of each cell and then subtracted the DAPI and LAMP-1 signals to define each cell’s cytoplasmic region (**Supplemental figure 2A**). We then measured the amount of p16 in each subcellular compartment by dividing the integrated pixel intensity of the p16 fluorescent signal by the number of pixels counted. This approach was used to normalize the concentration of p16 to the differing sizes of each compartment, as well as to the number of lysosomes present in each individual cell.

With our segmentation and quantification protocol in place, we tested the effect of inhibiting lysosomal degradation during stress on the expression and localization of p16. Targeting of p16 to lysosomes was first induced by treating RPE-1 cells with 20 µM etoposide. After 48 hours, cells were additionally treated with 40 mM NH_4_Cl for 24 hours to raise luminal pH and block degradation by lysosomes. In DMSO-treated cells, mean cytoplasmic, nuclear, and lysosomal-p16 increased in response to NH_4_Cl treatment (**Figure 3D**). Etoposide treatment alone increased the mean lysosomal p16 of the population relative to control, while overall mean cytoplasmic and nuclear p16 levels remained unchanged. However, although population mean was unchanged, etoposide-treatment bifurcated the population into relatively low and high p16 concentrations within those two compartments. Interestingly, in contrast to DMSO-treatment, when lysosomal degradation was inhibited by NH_4_Cl in etoposide-treated cells, lysosomal levels of p16 were unchanged. Instead, these cells were found to have significantly higher levels of nuclear p16.

In order to determine if p16 was preferentially targeted to the nucleus in response to lysosomal dysfunction during stress, we created a p16 “localization score” by dividing the mean nuclear, cytoplasmic, or lysosomal p16 fluorescence intensities by the mean p16 intensity of the entire cell (**Figure 3E**). This normalization procedure provides a stringent metric of p16 localization by detecting changes in p16 relative to any cell-wide changes in p16 levels, thereby allowing us to determine where the protein was preferentially recruited. Analyzing the p16 localization score in response to each treatment condition revealed that etoposide shifted preferential p16 accumulation to lysosomes and away from the nucleus and cytoplasm. Treatment with NH_4_Cl attenuated lysosomal localization in response to etoposide and instead shifted p16 accumulation to the nucleus. Furthermore, these cells developed a positive correlation between lysosomal and nuclear p16 localization in response to lysosomal inhibition that was not present in untreated cells (**Figure 3F**), suggesting the facilitation of p16 transport between these two subcellular components. In contrast, lysosomal inhibition did not change the correlation between cytoplasmic and nuclear p16 (**Supplemental figures 2B**), demonstrating specific interconnectivity between lysosomal and nuclear p16 localization. These results suggest that the combination of genotoxic stress and autophagy triggered by etoposide leads to competition between lysosomal and nuclear p16 enrichment.

### Inhibition of autophagy leads to accumulation of p16

The previous experiments showing that p16 localizes with lysosomes led us to ask if the autophagy pathway contributes to p16 degradation. Inhibiting autophagy results in the accumulation of p62 and other proteins destined for lysosomal degradation [21]. Here, we examined the effects of inhibiting autophagy on p16 protein levels using a variety of drugs to probe specific members of the autophagy pathway (**Figure 1**). As described previously, ammonium chloride prevents lysosomal degradation by raising luminal pH. Degradation can also be perturbed through direct inhibition of lysosome-associating vesicles. For example, bafilomycin A1 and chloroquine are potent inhibitors of late stage autophagy that act by preventing fusion between autophagosomes and lysosomes [28];[29].

RPE-1 cells were treated with autophagy/lysosome inhibitors bafilomycin A1 (BAFA1, 100 nM), chloroquine (CHQ, 20 µM), and ammonium chloride (NH_4_Cl, 40 mM), or the proteasome inhibitor MG132 (0.5 µM) for 24 h. Immunoblotting of whole cell lysates revealed increased p16 protein levels with either lysosome or proteasome inhibitors (**Figures 4A and 4B**). p62 levels also increased with every autophagy inhibitor, confirming dysregulation of the autophagosome-lysosome pathway. To determine if inhibition of autophagy promoted senescence, we also stained for phosphorylated-Rb on serine 780, a site specifically targeted by cyclin D1-CDK4 when the cell cycle is active [30]. Every autophagy inhibitor tested resulted in significantly lower phosphorylated-Rb (S780). RT-qPCR revealed that increases in p16 protein in response to autophagy inhibition were not the result of *de novo* p16 transcription (**Figure 4C**). Together these results demonstrate that p16 can be degraded by the autophagy/lysosomal pathway and implicate autophagy as a potential regulator of cell cycle progression.

**Figure 4.**
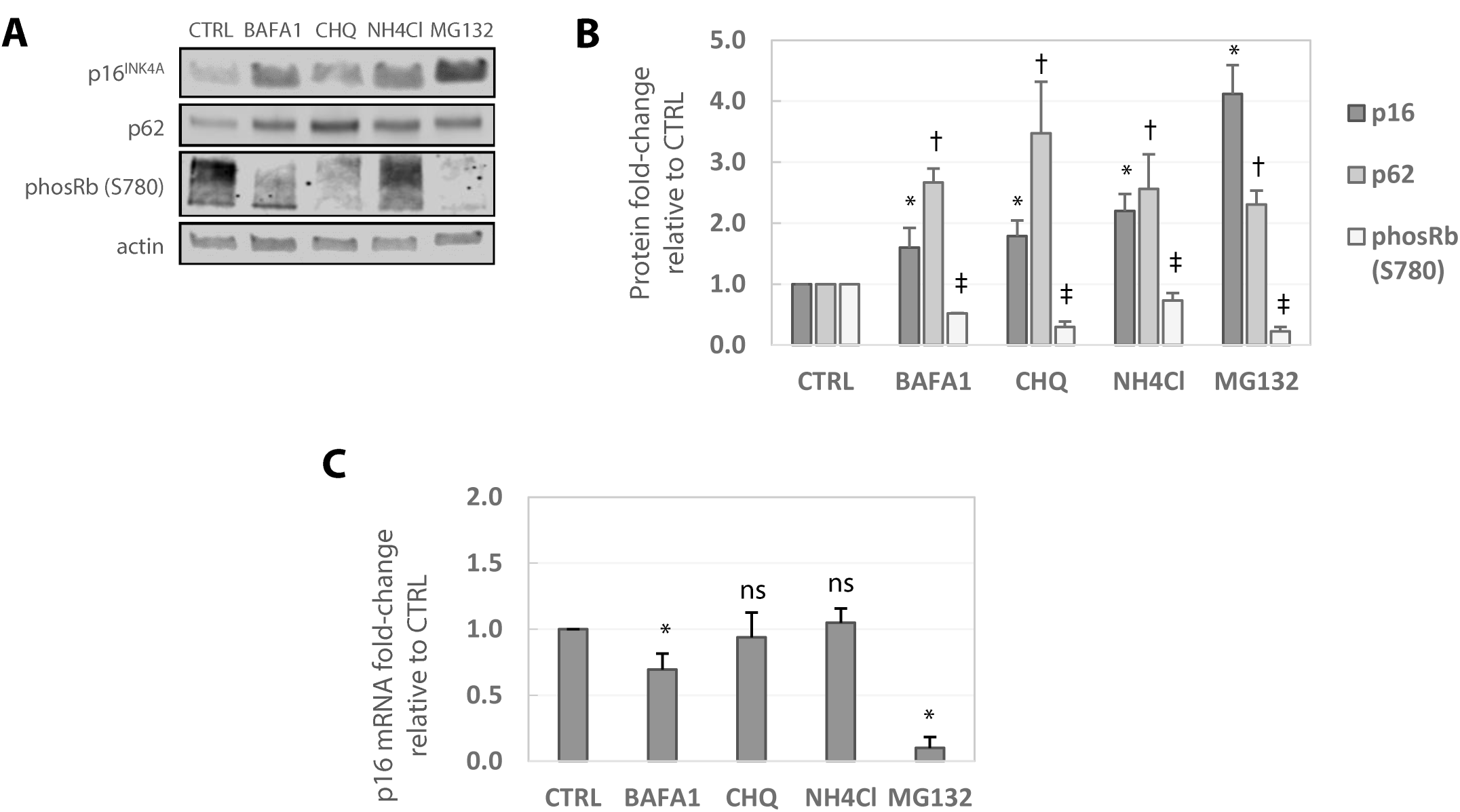
Blocking autophagy increases p16 protein but not transcript levels. RPE-1 cells were treated with autophagy/lysosome inhibitors bafilomycin A1 (BAFA1, 100 nM), chloroquine (CHQ, 20 μΜ), or ammonium chloride (NH_4_CI, 40 mM); proteasome inhibitor MG132 (0.5 μΜ); or DMSO vehicle control (CTRL) for 24 hours. **A)** Representative Western blot. **B)** Quantification of Western blot in panel A. *,†, ‡ = p < 0.05 relative to respective CTRL. **C)** RT-qPCR measuring p1 6 transcript levels. * = p < 0.05 relative to CTRL. ns = p > 0.05.

### p16 is targeted to autophagosomes by p62/SQSTM1

Proteins destined for lysosomal degradation are first targeted to autophagosomes and endosomes produced by the endomembrane system. We therefore asked if p16 is targeted to autophagosomes. Immunofluorescence revealed that autophagy inhibitor chloroquine perturbed the clearance of autophagosomes, resulting in stalled and enlarged cytoplasmic vacuoles containing autophagosome marker LC3 (**Supplemental figure 3A**). To determine if p16 is targeted to autophagosomes, we treated RPE-1 cells with DMSO or 20 μM chloroquine for 24 hours. Immunofluorescence revealed that DMSO-treated cells stained positive for diffuse cytoplasmic p16 and concentrated nuclear p16 (**Figure 4A and 4B**). In these cells, the autophagosome marker LC3 and chaperone p62 were either sparse or undetectable, a phenomenon known to be caused by rapid autophagic flux [18]. Disruption of autophagy via chloroquine treatment caused aggregation of cytoplasmic p16 puncta, which co-localized with LC3-and p62-positive puncta, indicating that p16 is targeted to autophagosomes (**Figure 4A and 4C**). Silencing of p62 via siRNA knockdown ablated p62 puncta and resulted in fewer cytoplasmic p16 aggregates forming in response to chloroquine treatment. LC3 puncta formation was unaffected by p62 knockdown. Additionally, knockdown of p62 combined with chloroquine resulted in delocalization of p16 puncta from LC3, indicating that p62 is responsible for the aggregation of p16 and its localization to autophagosomes. In contrast, siRNA knockdown of p16 eliminated p16 puncta without disrupting LC3 and p62 expression and co-localization, demonstrating that p16 is not necessary for autophagosome formation (**Supplemental figures 3B and 3C**). The dependence of p16 aggregation on p62 was further suggested by immunofluorescence staining of chloroquine-treated cells, which revealed a significant correlation between p62 and p16 fluorescence intensity co-localized within LC3 puncta (**Figure 4D**). Taken together, these results demonstrate that autophagy triggers localization of p16 to autophagosomes in a manner that is dependent on the expression of the chaperone p62.

## Discussion

In summary, our study demonstrates that localization and degradation of the p16 protein is regulated in part by the autophagy-lysosomal pathway. Lysosomal p16 enrichment occurs in response to DNA damage and autophagy induced by the chemotherapeutic agent etoposide. However, genotoxic stress in the presence of lysosome dysfunction leads to preferential nuclear accumulation of p16. Inhibiting autophagy raises p16 protein levels within 24 h without stimulating *de novo* p16 gene expression, demonstrating that p16 can be degraded by this pathway. Additionally, we found that recruitment of p16 to autophagosomes requires the chaperone protein p62. Together, these results reveal an unappreciated mode of regulation of the p16 protein in human cells.

Traditionally, protein localization has been studied with immunohistological experiments using antibodies targeting the protein of interest. However, these methods require the fixation of cells, which prevents temporal analysis of protein expression and localization. The autophagy pathway and endomembrane system is dynamic, mobile, and known to induce drastic changes in protein localization in a relatively short time-frame. By creating an endogenous p16-mCherry reporter in human cells, we have contributed a novel tool for examining p16 expression and translocation over time. Utilization of this reporter in future experiments will help to further our understanding of p16 dynamics in response to chemotherapeutic agents, cellular stress, and autophagy dysfunction.

Further study is required to identify the precise mechanisms that control p16 localization. For example, it is not known which domains on the p16 protein are responsible for autophagosomal and lysosomal recruitment. While we have found that p62 is required for p16 recruitment to autophagosomes, the endomembrane-transport system is complex, with many additional chaperone proteins and post-translational modifiers involved in recruiting, sorting, and shuttling cargo between different compartments of the cell. Determining the specific factors that control p16 transport could reveal potential drug targets for disease and anti-aging therapies.

We found that inhibiting autophagy led to p16 accumulation, decreased phosphorylated-Rb, and perturbed cell growth. Furthermore, the general lack of p16 aggregates under basal conditions implies that, in the cell lines tested, p16 remains in autophagic flux, which may allow replicating cells to sustain basal levels of p16 expression without inducing cell-cycle arrest. Additionally, we have demonstrated that activation of autophagy recruits p16 to lysosomes for degradation, which may prevent p16-induced senescence despite increases in p16 expression stimulated by cellular stress. It is intriguing, therefore, that inhibiting lysosomal degradation leads to nuclear accumulation of p16. Previous studies have shown that there is considerable cross-talk between lysosome-to-nuclear signaling and stress response proteins, including mTOR [27]. Our study expands this relationship to the tumor suppressor p16, and links p16 localization to lysosomal function, which both serve as key regulators of senescence, disease, and aging.

Since the p16 protein has long been known to promote cell cycle arrest through inhibition of CDK4/6 in the nucleus, these results suggest—but do not prove—a potential competition between the autophagy and senescence pathways through the sequestration of p16 (**Figure 5A**). Under this hypothetical model, stress induces the slow production of p16, which is quickly recruited to autophagosomes and degraded by lysosomes via the autophagy pathway. Over time, either through enhanced transcriptional activity or through p16 protein localization outside of lysosomes, p16 is able to enter the nucleus to bind to CDK4/6 and arrest the cell cycle. However, if the autophagy pathway is inhibited, p16 preferentially accumulates in the nucleus, which could lead to premature senescence (**Figure 5B**). From this model, we posit that autophagy “buys time” for cells undergoing stress to determine whether the damage is manageable and cells are able to resume proliferation once the stress conditions are eliminated. Alternatively, if stress conditions persist, or if autophagy is dysregulated, the cell enters senescence. Future studies will be necessary to determine whether sequestration of p16 through the autophagy-lysosomal pathways reduces a cell’s tendency to undergo senescence.

**Figure 5.**
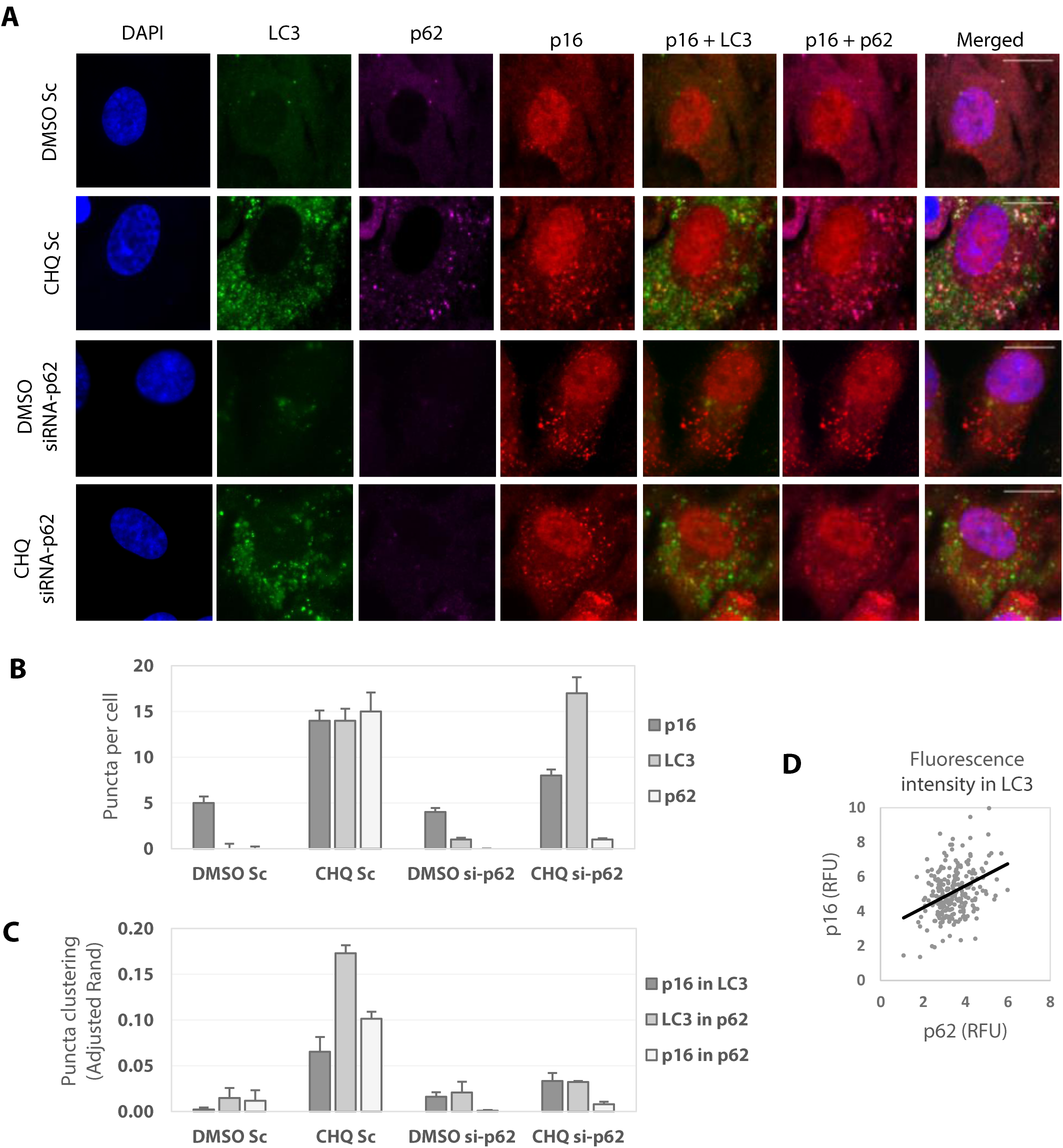
p16^INK4A^ is targeted to autophagosomes by p62. Human RPE-1 cells were treated with 20 μΜ chloroquine (CHQ), DMSO vehicle control, and/or transfected with siRNA-p62 or scramble control (Sc) for 24 hours. Cells were then fixed with paraformaldehyde, permeabilized with digitonin, and immunofluorescence stained. Scale bars = 15 μm. **A)** DMSO control cells stain positive for diffuse cytoplasmic p16 (red) and concentrated nuclear p16. Autophagasome markers LC3 (green) and p62 (magenta) are undetectable. Chloroquine treatment causes aggregation of cytoplasmic p16 puncta co-localized with LC3- and p62-positive puncta. Knockdown of p62 results in aggregation of cytoplasmic p16 puncta. p62 knockdown combined with chloroquine results in p16 puncta that do not co-localize with LC3-positive vesicles. **B)** Quantification of puncta segmented per cell. **C)** Quantification of puncta clustering using adjusted Rand index (see Materials and Methods). **D)** Regression plot of CHQ Sc sample comparing mean p16 fluorescence intensity versus mean p62 fluorescence intensity overlapping segmented LC3 puncta. RFU = mean pixel intensity x 1000. Pearson's correlation coefficient r = 0.35, df = 237, p-value = 4.8E-08.

**Figure 6.**
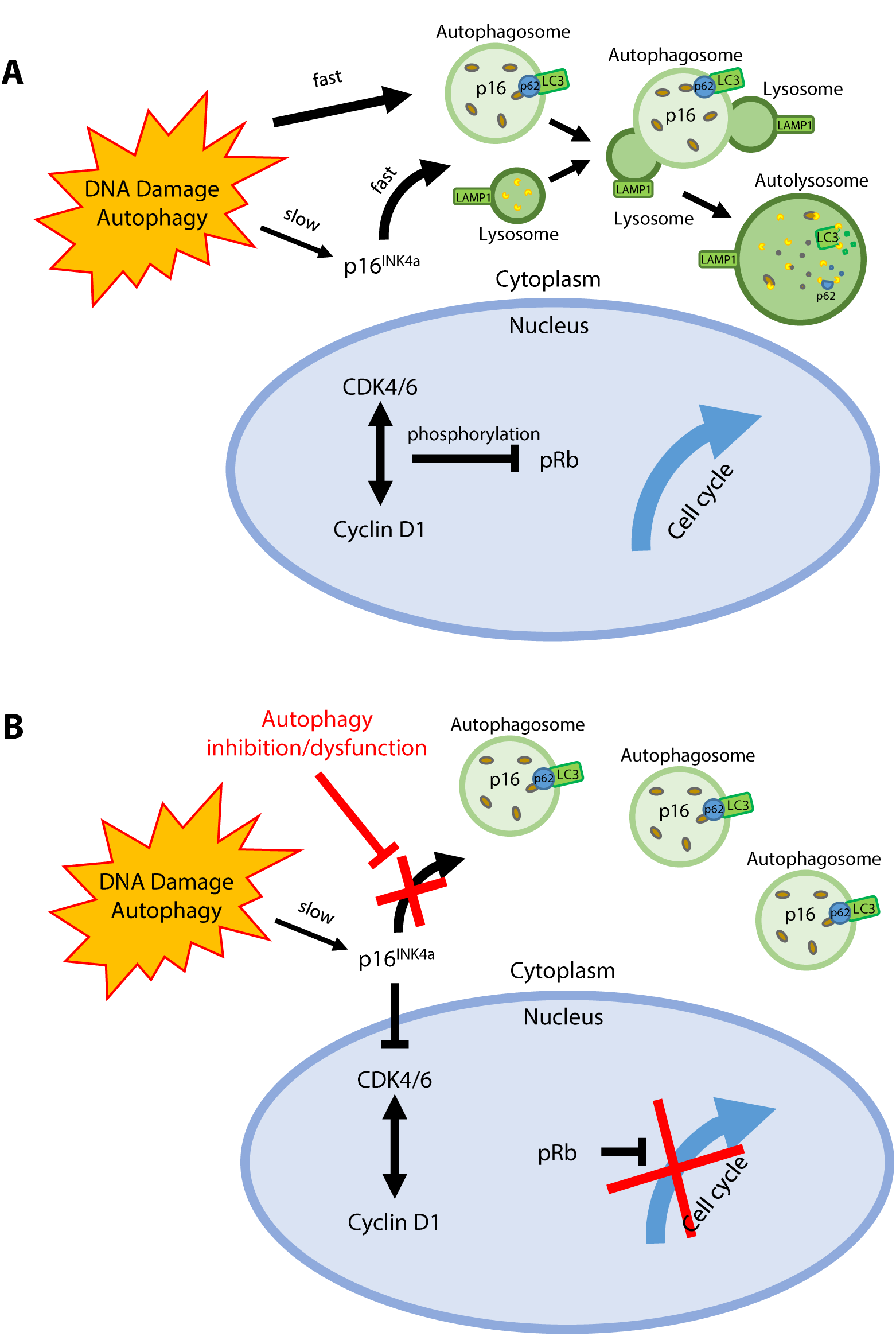
Hypothetical p16 autophagy-senescence competition model. **A)** In response to genotoxic stress, autophagy promotes rapid sequestration and degradation of p16 in the cytoplasm, preventing nuclear accumulation of the protein. Accumulation of p16 via de novo transcription is relatively slower and requires persistent stress. **B)** When autophagy-lysosome function is blocked, either by inhibiting vesicular trafficking or lowering pH, p16 is re-routed to the nucleus. Further work is necessary to determine whether sufficient p16 can accumulate under these conditions to induce cellular senescence.

Finally, the observation that p16 localizes to and is degraded by lysosomes represents a potentially novel thread of research for cancer cell biology. Commonly used chemotherapeutic drugs can induce increases in p16 expression in patients, but the effect of these agents on p16 localization in single cells has not been fully explored. Understanding how these compounds effect p16 localization could illuminate our understanding of how these treatments work at a mechanistic level. The ability to control senescence and attenuate cell growth via combined treatment with chemotherapeutics and well-established autophagy inhibitors could have major implications in the treatment of tumor progression. Beyond this application, the ability to slow or prevent senescence in healthy proliferating cells, such as stem cells, could lead to potential new therapies for other age-related diseases. In addition, we believe it is worth exploring the role of p16 in lysosomal storage diseases, which account for dozens of disorders associated with the brain, skin, heart, and central nervous system.

## Materials and Methods

### Cell culturing and maintenance

For routine maintenance and growth, RPE-1 cells were maintained in culture medium consisting of MEM (Gibco 11095080) supplemented with 10% FBS. For live-cell fluorescent microscopy experiments, RPE p16-mCherry cells were maintained in FluoroBrite culture medium consisting of DMEM (Gibco A1896701) supplemented with 10% FBS and 1% L-glutamine.

### Autophagy inhibitors

For autophagy inhibition experiments, cells were treated in culture medium supplemented with either NH_4_Cl (Sigma A9434), chloroquine (MedChemExpress HY-17589), bafilomycin A1 (MedChemExpress HY-100558), or MG132 (MedChemExpress HY-13259).

### Live-cell experiments and fluorescence quantification

RPE p16-mCherry reporter cells were plated onto glass-bottom 12-well plates in FluoroBrite culture medium. After 24 hours, media was replaced with FluoroBrite culture medium supplemented with either 20 µM etoposide (MedChemExpress HY-13629) or 0.5% DMSO alone, or in combination with 40 mM NH_4_Cl. Live-cell fluorescent microscopy was then performed for 24 hours.

Segmentation, counting, and fluorescence quantification of cells and subcellular compartments was performed in CellProfiler. Measured results were plotted and tested for statistical significance using MATLAB.

### Immunofluorescence, siRNA, and fluorescence quantification

RPE-1 cells were plated onto glass-bottom 12-well plates in culture medium. After 24 hours, media was replaced with culture medium supplemented with either 20 µM etoposide or 0.5% DMSO. Depending on experimental conditions, cells were also treated with Lipofectamine RNAiMAX Transfection Reagent (ThermoFisher 13778030) and siRNA-p16 (Dharmacon), siRNA-p62 (Dharmacon), or non-targeting scramble siRNA (Dharmacon D-001206-13-05). 48 hours after treatment, media was inoculated with either 40 mM NH_4_Cl, 20 µM chloroquine, or left untreated for 24 hours. Cells were washed with ice cold PBS and fixed with 4% PFA for 10 minutes at room temperature. Cells were permeabilized and blocked with 0.02% digitonin (Invitrogen BN2006) in PBS containing 5% serum for one hour at room temperature. All following steps were performed in wash buffer containing 5% serum in PBS. First, cells were incubated overnight at 4°C in wash buffer containing Anti-CDKN2A/p16INK4a antibody (Abcam ab108349), Anti-LAMP1 antibody (Abcam ab25630), Anti-SQSTM1/p62 antibody (Abcam ab56416), or Anti-LC3B antibody (Abcam ab192890). The following day, cells were washed three times for five minutes with wash buffer. Cells were then incubated for one hour at room temperature in wash buffer containing mouse and rabbit conjugated secondary antibodies. For p62 and LC3 staining experiments, cells were then washed three times for five minutes with PBS, blocked again for one hour in wash buffer, and then incubated overnight at 4°C in wash buffer containing Anti-CDKN2A/p16INK4a conjugated (Alexa Fluor 647) antibody (Abcam ab192054). Cells were then washed with PBS containing 1 µg/mL DAPI for five minutes at room temperature, followed by three five-minute washes with PBS before visualization.

Segmentation, counting, and fluorescence quantification of cells, subcellular compartments, and fluorescent puncta were performed in CellProfiler. Puncta clustering analysis was performed in CellProfiler using the adjusted Rand index measured between fluorescent channels. Measured results were plotted and tested for statistical significance in MATLAB using ANOVA and Bonferroni correction for multiple comparison analysis.

### Western blot and protein quantification

RPE-1 cells were plated onto Corning 6-well cell culture plates in culture medium. The next day, media was replaced with culture medium supplemented with either 40 mM NH_4_Cl, 20 µM chloroquine, 100 nM bafilomycin A1, 0.5 µM MG132, or 0.5% DMSO for 24 hours. For whole-cell protein analysis, cells were lysed with ice cold RIPA buffer containing protease and phosphatase inhibitors. Lysates were separated on a gradient gel (TGX, BioRad), and transferred to a PVDF membrane. Membranes were blocked with blocking buffer (LI-COR Odyssey Blocking Buffer 927-40000) for 1 hour before probing with primary antibodies for p16 (Abcam ab108349), SQSTM1/p62 (Abcam ab56416), phospho-Rb S780, and beta-actin (sigma) in blocking buffer overnight at 4°C. Membranes were washed and probed with secondary antibodies (LI-COR IRDye800 and IRDye680) for 1 hour at room temperature and visualized using the LI-COR Odyssey CLx Imaging System. Proteins were normalized to actin and quantified using ImageJ.

## Acknowledgements

We thank Samuel Wolff for guidance on fluorescence microscopy; Kasia Kedziora for assistance with image analysis; Brian Diekman for helpful discussions about p16 and senescence; Norman Sharpless for assistance in project conceptualization and valuable feedback; and members of the Purvis Lab for technical suggestions and support. This work was supported by the National Institutes of Health research grants T32-GM007092D (P.C.), R00-GM102372 (J.E.P.), and DP2-HD091800 (J.E.P.); a Medical Research Grant from the W.M. Keck Foundation (J.E.P.); and the North Carolina University Cancer Research Fund.

## Author Contributions

P.R.C. conceptualized the project and all experiments, constructed the RPE-1 p16-mCherry 53bp1-mVenus reporter cell line, performed image analysis, created figures, and wrote the manuscript. P.R.C., S.K.G., and K.A.G. performed cell maintenance, Western blot, immunofluorescence, and qPCR experiments. M.A.R. assisted in cell maintenance and Western blot experiments. S.R.S. assisted in cell maintenance and immunofluorescence experiments.

---

**Supplemental figure 1.**
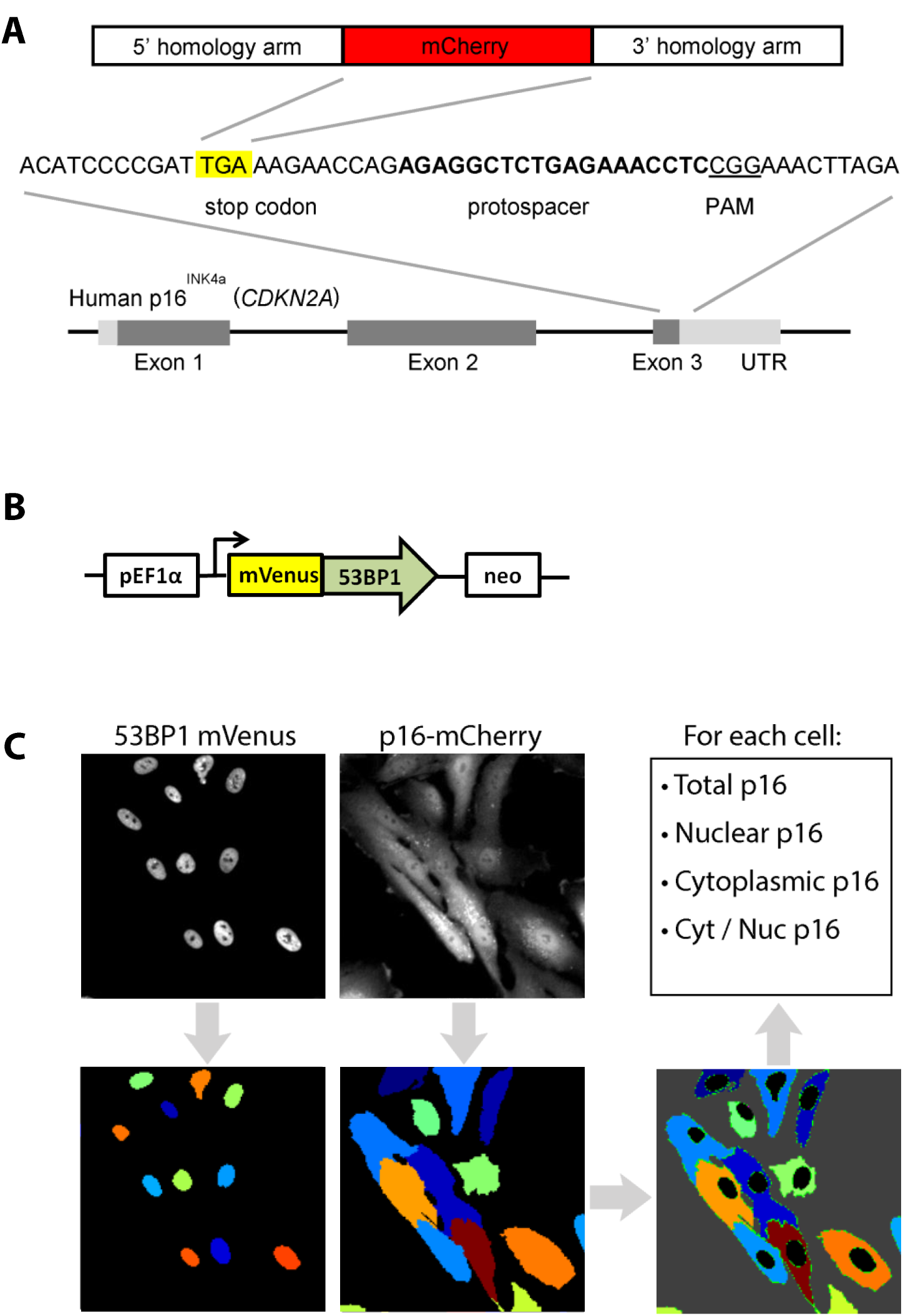
Development of a p16-mCherry 53bp1 -mVenus reporter cell line. **A)** Schematic for CRISPR/Cas9-mediated mCherry incorporation at the p1 6/CDKN2A stop codon. **B)** Schematic for constitutive 53bp1 -mVenus cassette incorporated into p16-mCherry cells via lentiviral transduction. **C)** 53bp1 -mVenus was used to identify and segment individual nuclei. Whole cellular area was identified by segmenting p16-mCherry signal throughout both the nucleus and cytoplasm. Cytoplasmic segmentation was performed by masking nuclei with their corresponding whole-cell.

**Supplemental figure 2.**
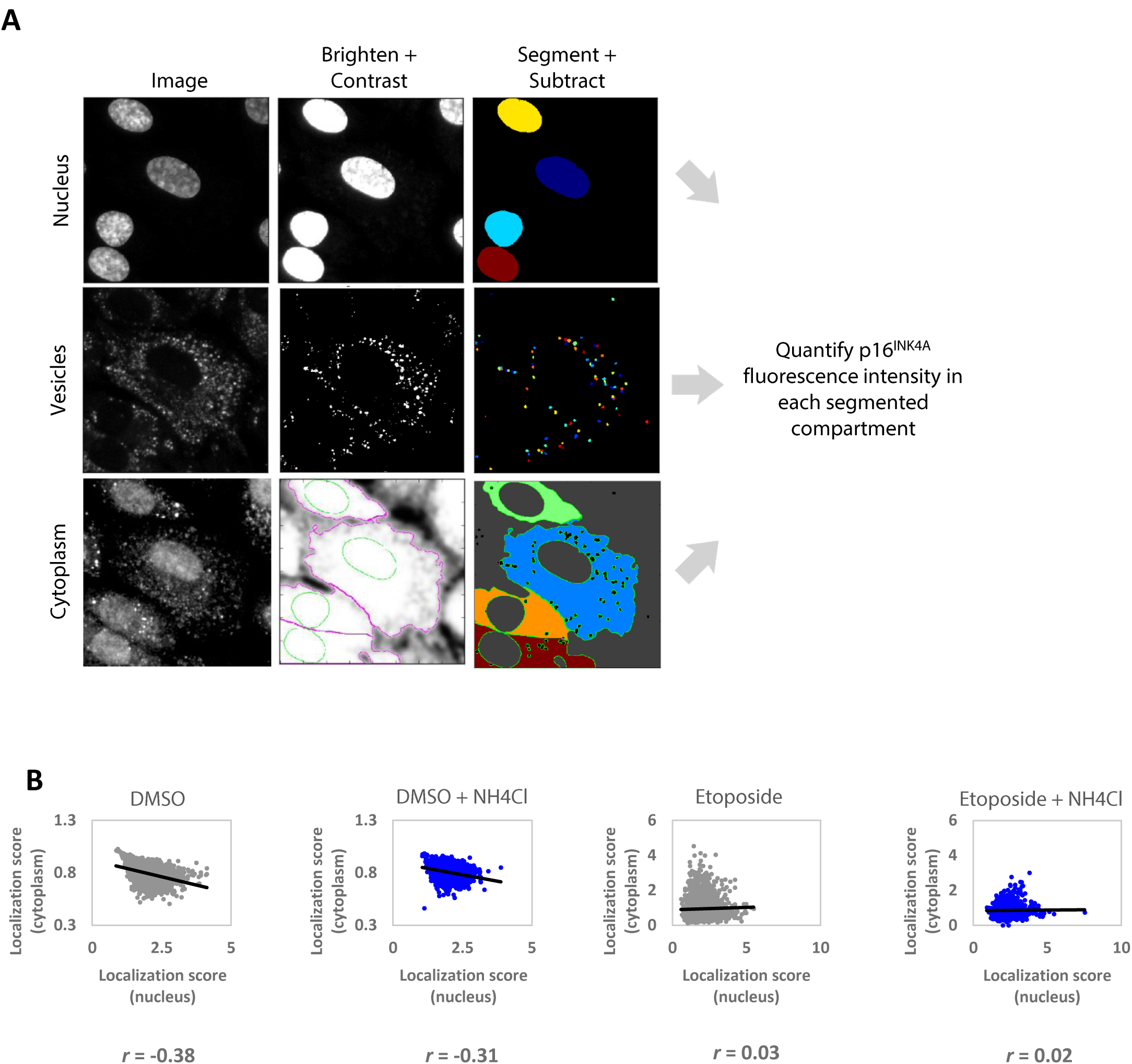
**A)** Subcellular segmentation of immunofluorescence images. Nuclear segmentation is performed using an image of cells stained with DAPI. Vesicle segmentation is performed by segmenting puncta formed by a vesicle-targeting antibody (LAMP-1, LC3, p62, etc.). Cytoplasmic segmentation is done by first segmenting whole cells by performing a watershed segmentation of brightened p16 images using the previously obtained nuclei. Nuclei and vesicles are then subtracted from the image. Each overlapping nucleus, vesicle, and cytoplasm is then grouped for single-cell analysis. p16 fluorescence intensity can then be quantified in each subcellular compartment (nuclear, vesicular, and cytoplasmic) for each cell. **B)** Regression plots comparing p16 cytoplasmic localization score vs. p16 nuclear localization score. *r* = Pearson's correlation coefficient.

**Supplemental figure 3.**
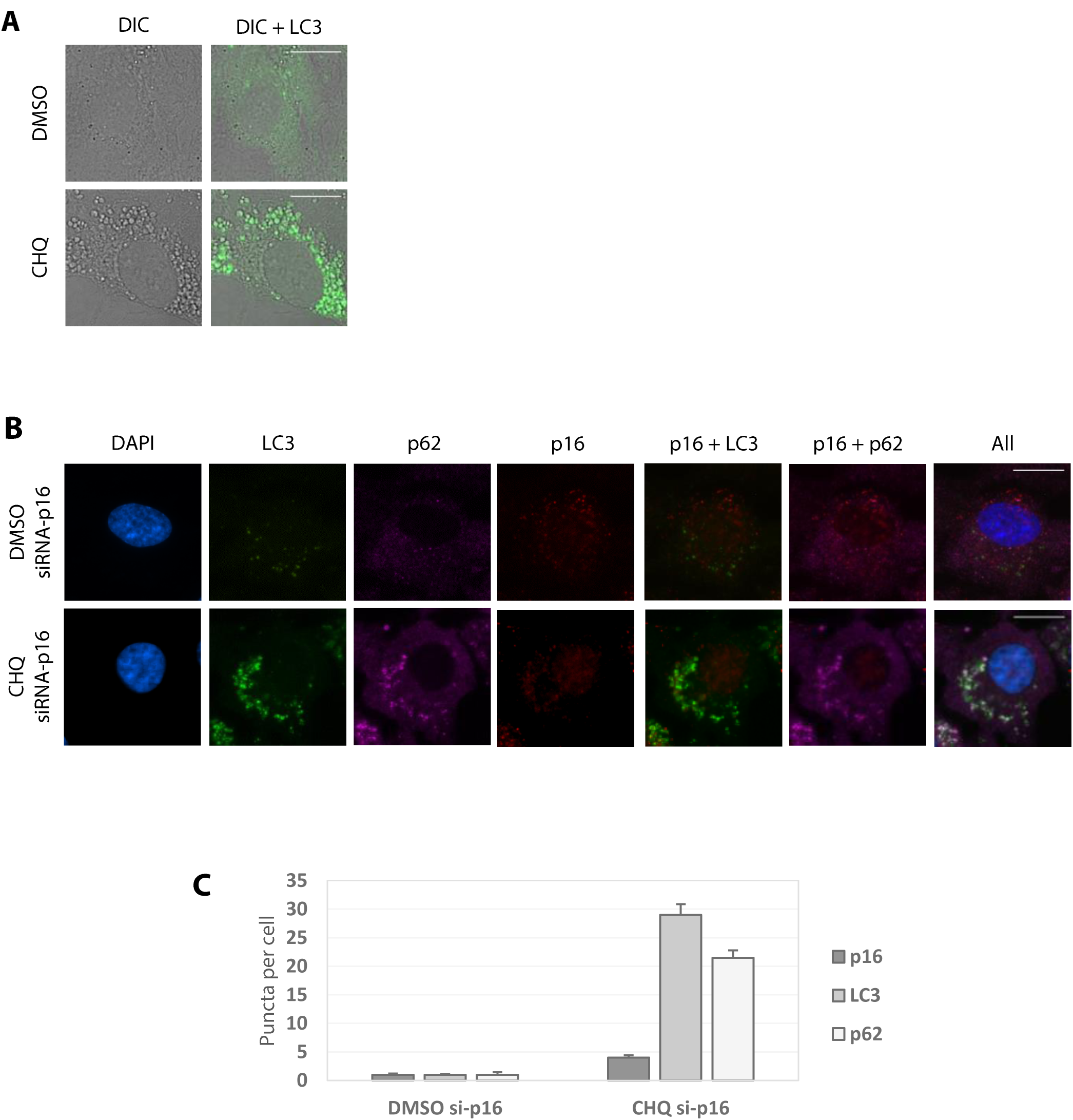
**A)** Human RPE-1 cells were treated with 20 μΜ chloroquine (CHQ) or DMSO vehicle control for 24 hours. CHQ forms enlarged cytoplasmic vesicles that stain positive for autophagasome membrane protein LC3 (green). **B)** Human RPE-1 cells were treated with 20 μΜ CHQ, DMSO vehicle control, and transfected with siRNA-p16 for 24 hours. Cells were then fixed with paraformaldehyde, permeabilized with digitonin, and immunofluorescence stained. Scale bars = 15 μm. DMSO control cells with siRNA-p16 stained negative for p16 (red) and autophagasome markers LC3 (green) and p62 (magenta). Chloroquine treatment causes aggregation of LC3- and p62-positive puncta, but not p16**. C)** Quantification of puncta segmented per cell.

